# Modulation of gamma spectral amplitude and connectivity during reaching predicts peak velocity and movement duration

**DOI:** 10.1101/2021.11.11.468153

**Authors:** Elisa Tatti, Francesca Ferraioli, Alberto Cacciola, Cameron Chan, Angelo Quartarone, M. Felice Ghilardi

**Author notes:** Corresponding authors: Elisa Tatti, CUNY School of Medicine, 160 Convent Avenue, Harris Hall room 008, New York, NY, 10031,; M. Felice Ghilardi, CUNY School of Medicine, 160 Convent Avenue, Harris Hall room 008, New York, NY, 10031.

## Abstract

Voluntary movements are accompanied by increased oscillatory activity or synchronization in the gamma range (> 25.5 Hz) within the sensorimotor system. Despite the extensive literature about movement-related gamma synchronization, the specific role of gamma oscillations for movement control is still debated. In this study, we characterized movement-related gamma oscillatory dynamics and its relationship with movement characteristics based on 256-channels EEG recordings in 64 healthy subjects while performing fast and uncorrected reaching movements to targets located at three distances. We found that movement-related gamma synchronization occurred during both movement planning and execution, albeit with different gamma peak frequencies and topographies. Also, the amplitude of gamma synchronization in both planning and execution increased with target distance. Additional analysis of phase coherence revealed a gamma-coordinated long-range network involving occipital, frontal and central regions during movement execution. Gamma synchronization amplitude and phase coherence pattern reliably predicted peak velocity amplitude and timing, thus suggesting that cortical gamma oscillations play a significant role in the selection of appropriate kinematic parameters during planning and in their implementation during movement execution.

## Introduction

Voluntary movements are accompanied by modulation of oscillatory activity of both the beta (13.5-25 Hz) and gamma frequency ranges (25.5-90 Hz) that have been consistently observed with EEG, MEG and ECoG over the sensorimotor cortex and with electrode recordings in basal ganglia structures.

Differently from event-related beta desynchronization (ERD) and synchronization (ERS) dynamics that can be elicited by passive movements and are not correlated with specific kinematic features (Cremoux et al., 2013; Pistohl et al., 2012; Salmelin et al., 1995; Stancák and Pfurtscheller, 1996, 1995; Tatti et al., 2019), gamma oscillations display a more direct association with the motor output. The notion of a prokinetic nature of movement-related gamma oscillations is supported by studies showing some relationship with characteristics of the movement, such as distance, duration, velocity and applied force level (Ball et al., 2008; Brücke et al., 2012; Cheyne and Ferrari, 2013; Gunduz et al., 2016; Joundi et al., 2012; Lofredi et al., 2018; Muthukumaraswamy, 2010; Nowak et al., 2018). Nonetheless, their specific role in movement planning and execution is still a matter of debate. Based on studies of gamma ERS in the primary somatosensory cortex following tactile stimulation (Christensen et al., 2007), gamma oscillations have been generally associated with the proprioceptive feedback ensuing movement and sensorimotor input. However, this hypothesis has been challenged by reports showing that passive movements do not elicit gamma synchronization (Brücke et al., 2012; Muthukumaraswamy, 2010) and that mirror illusion in absence of proprioceptive feedback prompts movement-related gamma ERS (Butorina et al., 2014). Additionally, although gamma ERS is prominent during movement execution, a few studies have reported increased gamma activity even before movement onset (Gaetz et al., 2013; Gunduz et al., 2016), thus suggesting that gamma oscillations might represent a signature of processes linked to goal-directed movement representation, planning and execution.

In recent studies, we have extensively used a reaching task where fast and uncorrected movements to different target distances result from a more prominent scaling of peak velocity rather than of movement duration (Tatti et al., 2019, 2020, 2021). With that task, we have demonstrated that the movement-related beta ERD-ERS dynamics does not depend on target distance, movement length or peak velocity (Tatti et al., 2019). In the present study, we used the same reaching task to investigate whether the amplitude of movement-related gamma ERS scales with movement distance and peak velocity. We thus recorded EEG activity in healthy young subjects performing reaching movements toward targets at three different distances and characterized changes of gamma oscillatory amplitude and phase-coherence activity. According to the existing evidence on the pro-kinetic nature of movement-related gamma ERS, we expected that, compared to shorter movements, longer movements would be associated not only with higher peak velocities but also with greater gamma ERS (Ball et al., 2008; Brücke et al., 2012; Cheyne and Ferrari, 2013; Gunduz et al., 2016; Lofredi et al., 2018; Muthukumaraswamy, 2010; Nowak et al., 2018; J. Wang et al., 2017). Further, we hypothesized that the amplitude and phase-synchronization of gamma oscillations could reliably predict such kinematic parameters.

## Results

### Movement characteristics depend on target distance

Sixty-four subjects performed 96 fast and uncorrected out and back movements from a central starting point to one of 24 targets (three distances and eight directions, Figure 1a) appearing in a random order. In general, the resulting movements were straight and their temporal velocity profiles were on average bell-shaped and appropriately scaled to the target distance, as displayed in Figure 1a and in previous publications (Tatti et al., 2019, 2020, 2021). Repeated measure ANOVAs showed that movement extent, peak velocity amplitude and timing, as well as movement duration increased with target distance (Supplementary Table 1, Figure 1b). Post-hoc tests revealed significant differences between the three target extents for all these measures. Target extent had a significant effect also on reaction time (Supplementary Table 1), with shorter reaction time for more distant targets. However, post-hoc analyses showed significant differences only between short targets and the other two targets.

**Figure 1.**
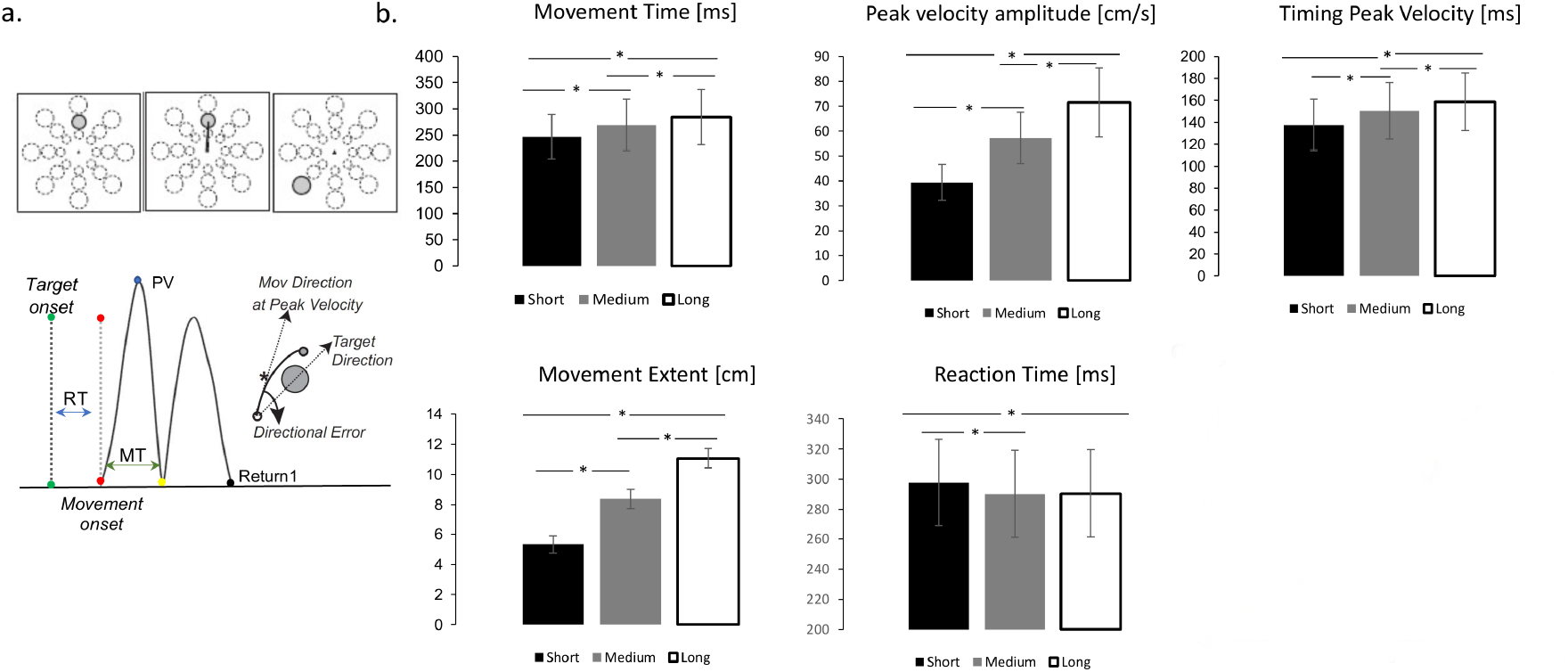
*mov* test depiction, kinematic indexes and performance. a) Top. mov test. One of 24 targets appeared in unpredictable order every 3 seconds. Bottom. Measures of movement characteristics. b. Mean and standard error for each target distance (Black= short, Gray= medium, White=Long) for the kinematic parameters. The horizontal lines identify significant pairwise comparisons after Bonferroni correction (p<0.05).

We then performed linear mixed-effect regression modeling to characterize the contribution of peak velocity amplitude and movement time to movement extent. Thus, we first assessed: adjusted R^2^, Bayesian Information Criterion (BIC, an index used in Bayesian statistics to select among two or more models), and the Theoretical Likelihood Ratio Test (TLRT, commonly used to compare the goodness of fit of two statistical models) (see Methods). Following the indication provided by these indices, the model with random intercept and slope showed the best fitting of the data (Supplementary Table 2). The results showed that both peak velocity and movement time were strong predictors of movement extent variability: the model with random intercept and slope for each participant explained approximately 95% of the movement extent variance (R^2^ adjusted=0.95, Supplementary Table 2). The major contributor to movement extent variability was the amplitude of peak velocity (standardized fixed slope: 1.06; CI: 1.01-1.10; t_(1,4743)_ =46.83; p < 0.0001), while the effect of movement time was less prominent (standardized fixed slope: 0.70; CI: 0.65-0.75; t_(1, 4743)_ =30.08; p < 0.0001).

Altogether, these findings show that movement extent resulted mainly from the scaling of the force to the appropriate target distance with a lesser contribution of movement duration.

### Movement-related gamma oscillatory dynamics

We next investigated the progression and the topographical distribution of gamma oscillatory activity (25.5-80 Hz) during movement planning and execution. We thus compared gamma oscillatory activity of each 24 ms-time bins to the average spectral power of the entire epoch (from 500 ms before the movement onset to 2400 ms after) with non-parametric Monte Carlo permutation testing. This analysis revealed significant electrode clusters in two time-windows: one from 152 to 52 ms before movement onset, that is, during the reaction time period (Cluster t=907.27, p=0.0003, CI: 0.0004) and the other from 52 to 500 ms after moment onset, that is, during the movement execution time (Cluster t= 13.120, CI:19.598, p=0.0001). As shown in Figure 2, before movement onset, gamma oscillatory activity increased in a cluster of electrodes over the centro-parietal region; during movement execution, the increase in gamma activity at first, involved electrodes over the occipital region, then spread to most of the scalp electrodes and ultimately returned to a centro-parietal cluster.

**Figure 2.**
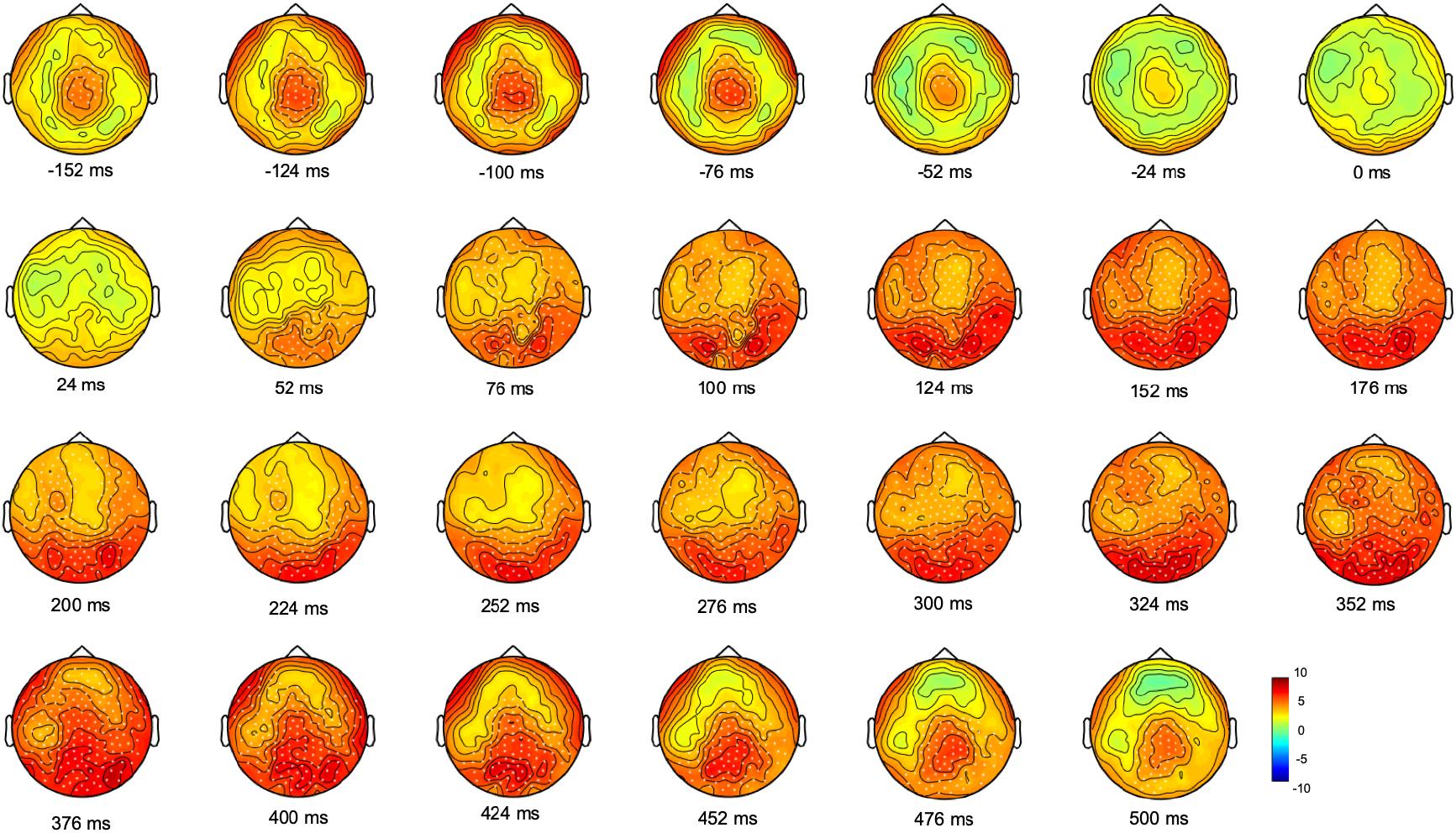
Monte Carlo cluster-based permutation T-statistic of the difference between gamma oscillatory activity and the average spectral power during movement planning and execution. Dots indicate significant clusters of electrodes (p<=0.0005).

We then averaged the topography of gamma activity in the time bins of the reaction time period (from 152 to 52 ms before movement) and that in the time bins from 52 to 500 ms after moment onset, during the movement execution time. Cluster-based permutation analyses on four gamma bands (broad gamma: 25.5-80 Hz, low gamma: 25.5-40 Hz, medium gamma: 40.5-55 Hz, high gamma: 55.5-80 Hz) revealed for both time windows significant cluster of electrodes in the broad, medium and high gamma bands (Figure 3). In all these gamma bands, the planning window displayed greater synchronization over a centro-parietal region, whereas movement execution showed a widespread gamma synchronicity over all scalp channels, with maximum amplitude increase over the occipital region. No significant clusters were found in the low gamma range. These results prompt some considerations about the role of gamma activity in both movement planning and execution. In line with a few reports (Gaetz et al., 2013; Gunduz et al., 2016), the occurrence of gamma synchronization over a centro-parietal region during movement planning suggests that gamma oscillations may reflect the generation of a motor output. In addition to the creation of a motor plan, gamma ERS might also be involved in the control of the actual movement, as suggested by the presence of gamma ERS increase during the motor act, thus indicating the possible engagement of online feedback control processes.

**Figure 3.**
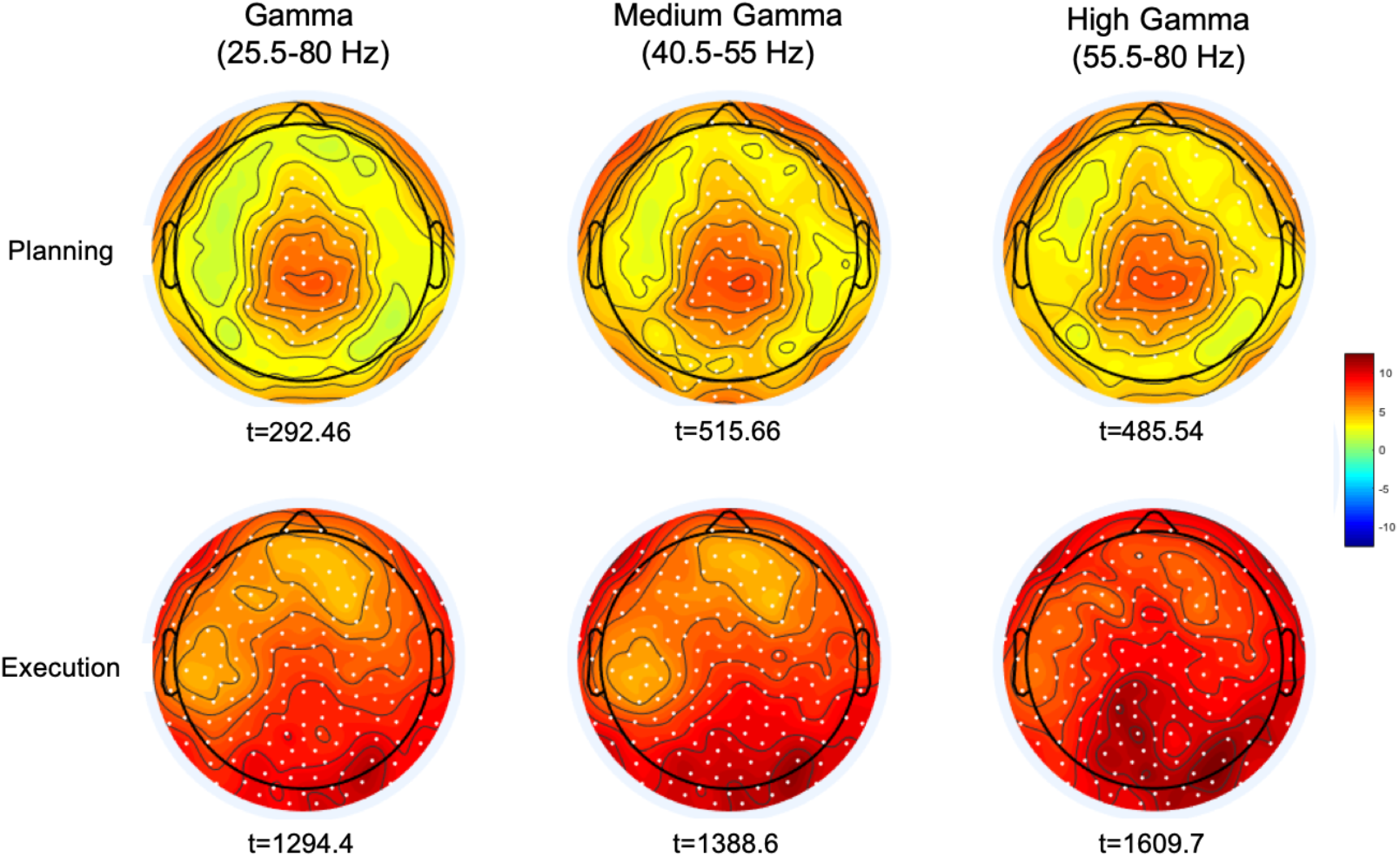
Maps of the cluster-corrected permutation-based t-values comparing gamma oscillatory activity with the average spectral power during the planning (−152 to −52 ms) and execution time-windows (52 to 500 ms). Significant electrodes are reported as white dots (p<0.0005).

### Movement-related gamma ERS shifts toward higher frequencies during movement execution

We then determined whether the observed gamma ERS during planning and execution differed in their peak frequency ranges (see Methods). We used non-parametric related-samples Friedman’s ANOVA to test possible differences of gamma ERS frequency between the planning, execution and post-movement time windows. As displayed in Figure 4, we found that gamma ERS peaked at different frequency ranges ( χ^2 (2)=^ 93.23, p<0.0001). During the planning window, gamma ERS was maximally expressed in the medium gamma range (53.73 Hz ±10.04, mean ± SD) whereas during movement execution the peak shifted towards higher frequencies (65.8 Hz ±10.72). Importantly, after movement completion, cortical activity synchronized back to the beta/low gamma frequency range (28.06 Hz ± 11.47; Dunn-Bonferroni post-hoc tests, Planning vs Movement: z = −0.594, p=0.002; Post-movement vs Planning: z = 1.68 p<0.0001; Post-movement vs Movement: z =1.086 p<0.0001). These findings demonstrate that gamma ERS occurring during movement planning and execution is maximally expressed in two distinctive frequency ranges, thus suggesting possible different functional properties of gamma ERS.

**Figure 4.**
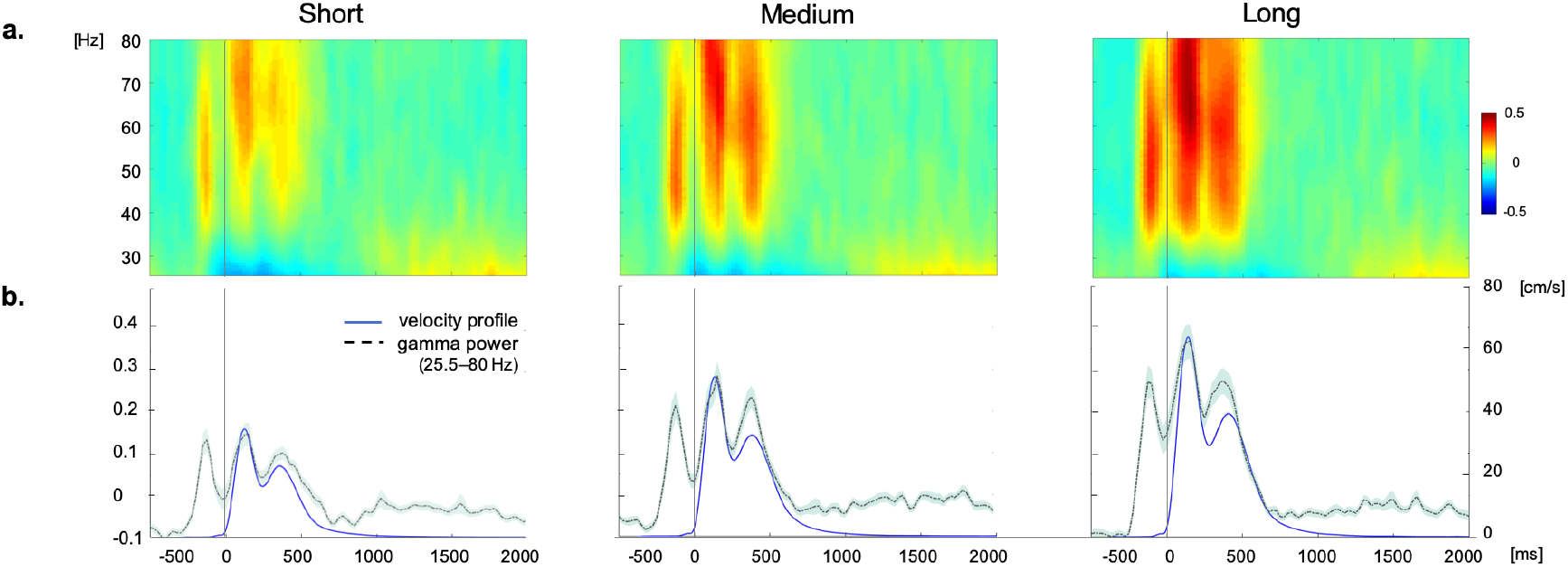
a) EEG time-frequency plots of the gamma band range (25.5-80 Hz) during movement separately averaged for target displayed at short (4 cm), medium (7 cm) and long (10 cm) distances. b) Time course of gamma power (black and velocity profile averaged across subjects for each target distance. Shaded area in the power and velocity profiles represents the standard error.

### Movement execution is characterized by greater long-range connectivity than planning

To further explore gamma activity differences between planning and execution, we compared their degree of gamma phase coupling across scalp channels. Phase-synchronization in the broad gamma and high gamma frequency range was measured with the weighted phase lag index (wPLI), a functional connectivity metric that minimizes the impact of volume conduction effects (see methods). The wPLI was computed across all channel-pairs to provide a whole-brain mapping of functional EEG networks and to ascertain possible differences of connectivity between the planning and the execution windows. Network-based statistics (NBS) revealed a sub-network of increased functional connectivity during movement execution compared to movement planning in the broad gamma frequency. The sub-network consisted of 244 edges connecting 92 different electrodes (p<0.001, corrected for multiple comparison). Interestingly, apart from a few pathways linking frontal to occipital regions, these patterns of increased connectivity mainly involved a left fronto-temporal-parietal network. (Figure 5a). No significant sub-networks of increased connectivity during movement planning were detected compared to the execution window. Since movement-related gamma ERS shifted towards a higher frequency range during movement execution, we thus looked for possible sub-network differences in the high-gamma range (55.5-80 Hz). Indeed, we found a sub-network of increased connectivity during movement execution compared to planning (p<0.001, corrected for multiple comparison) that consisted of 571 edges connecting 122 different electrodes. In addition to patterns of connectivity mainly involving the fronto-temporal-parietal areas bilaterally, this sub-network also involved fronto-occipital connections (Figure 5b), in line with our finding of greater high-gamma power increase over the occipital region during movement execution.

**Figure 5.**
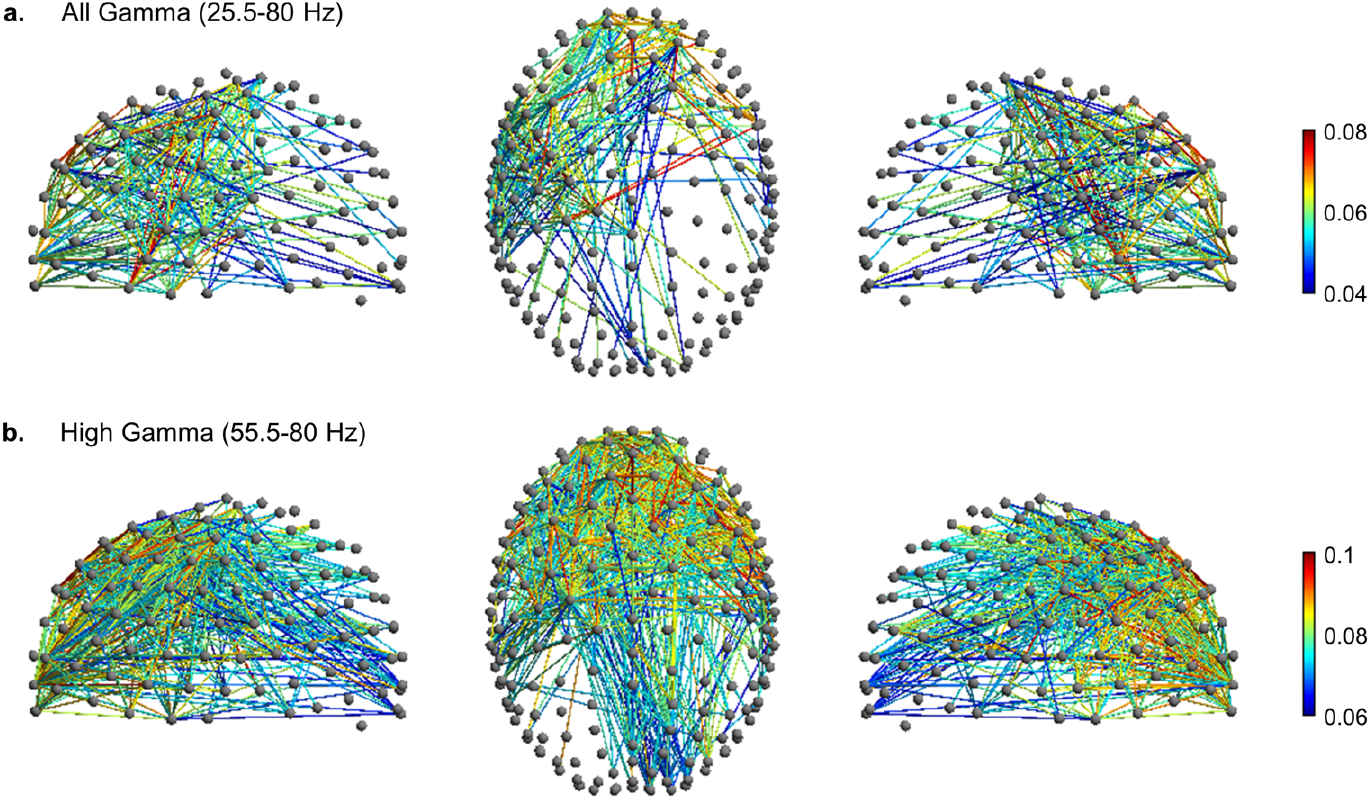
Movement-related significant sub-network with increased connectivity in the movement execution compared to the planning phase (p<0.001, corrected for multiple comparison) in the a. broad gamma (25.5-80 Hz) and b. high gamma (55.5-80 Hz) frequency band. The nodes and the links are depicted in two different projections (sagittal, on the left and right side; axial, in the center). Colormap indicates the difference of the mean of wPLI values between the movement execution and planning.

### Movement-related gamma ERS is modulated by target distance

We then investigated possible relationships between gamma activity indices and movement features. We first ascertained whether target distance would affect the amplitude of gamma ERS by comparing gamma oscillatory activity for short, medium and long target distances with cluster-based permutation analyses based on the significant two temporal windows, two electrode clusters and gamma bands identified in the previous analyses (Figure 3).

Both the omnibus repeated measure ANOVA and the subsequent post-hoc tests confirmed that, during both movement planning and execution, broad gamma oscillatory activity increased with target distance (Table 1a) with significant differences between short and medium and short and long target distance trials (Table 1b; Figure 4). Despite greater gamma power for long than medium distances in both planning (mean difference ± SD: 8.1%±18.4%) and movement execution (7.9%±13.0%), post-hoc tests did not reveal any significant differences between medium and long-distance targets. Similar results were obtained for medium gamma and high gamma frequency bands, except for medium and short movement trials during planning in the medium gamma frequency band (Table 1b).

**Table 1.**
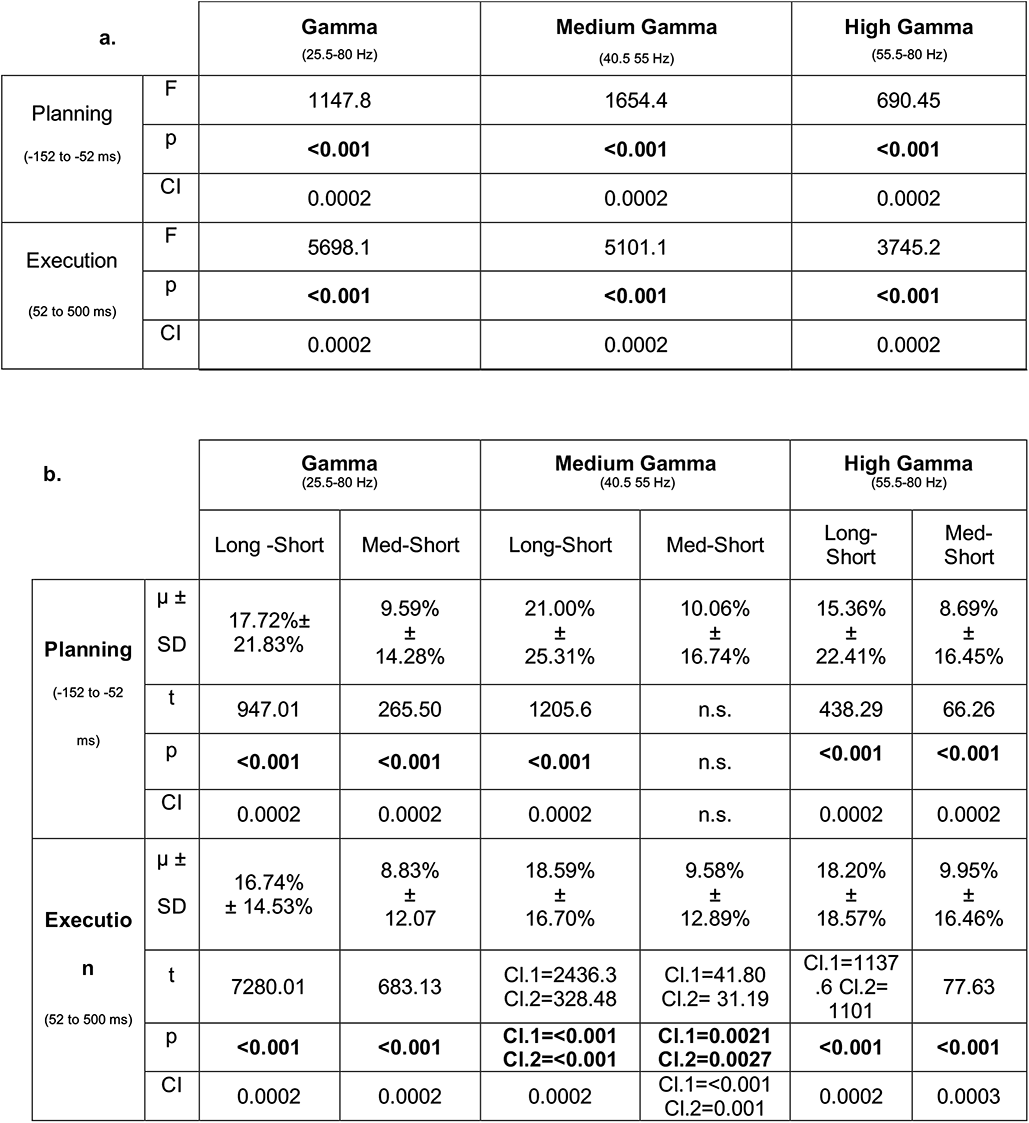
Non-parametric Monte Carlo permutation analysis F statistics and post-hoc tests with cluster correction on the difference in gamma amplitude between Short, Medium and Long target distance trials. μ= mean power difference; SD: Standard Deviation of the mean; CI: Confidence Interval, n.s.: not significant. Significant results are reported in bold.

| **Table 1 a.** |  | **Gamma**<br>(25.5-80 Hz) | **Medium Gamma**<br>(40.5 55 Hz) | **High Gamma**<br>(55.5-80 Hz) |
| --- | --- | --- | --- | --- |
| Planning<br>(-152 to -52 ms) | F | 1147.8 | 1654.4 | 690.45 |
|  | p | **<0.001** | **<0.001** | **<0.001** |
|  | CI | 0.0002 | 0.0002 | 0.0002 |
| Execution<br>(52 to 500 ms) | F | 5698.1 | 5101.1 | 3745.2 |
|  | p | **<0.001** | **<0.001** | **<0.001** |
|  | CI | 0.0002 | 0.0002 | 0.0002 |

**Table 1.** Non-parametric Monte Carlo permutation analysis F statistics and post-hoc tests with cluster correction on the difference in gamma amplitude between Short, Medium and Long target distance trials.
| **b.** |  | **Gamma**<br>(25.5-80 Hz) |  | **Medium Gamma**<br>(40.5 55 Hz) |  | **High Gamma**<br>(55.5-80 Hz) |  |
| --- | --- | --- | --- | --- | --- | --- | --- |
|  |  | Long -Short | Med-Short | Long-Short | Med-Short | Long-Short | Med-Short |
| Planning<br>(-152 to -52 ms) | $\mu \pm$<br>SD | 17.72%±<br>21.83% | 9.59%<br>±<br>14.28% | 21.00%<br>±<br>25.31% | 10.06%<br>±<br>16.74% | 15.36%<br>±<br>22.41% | 8.69%<br>±<br>16.45% |
|  | t | 947.01 | 265.50 | 1205.6 | n.s. | 438.29 | 66.26 |
|  | p | **<0.001** | **<0.001** | **<0.001** | n.s. | **<0.001** | **<0.001** |
|  | CI | 0.0002 | 0.0002 | 0.0002 | n.s. | 0.0002 | 0.0002 |
| Executio<br>n<br>(52 to 500 ms) | $\mu \pm$<br>SD | 16.74%<br>± 14.53% | 8.83%<br>±<br>12.07 | 18.59%<br>±<br>16.70% | 9.58%<br>±<br>12.89% | 18.20%<br>±<br>18.57% | 9.95%<br>±<br>16.46% |
|  | t | 7280.01 | 683.13 | Cl.1=2436.3<br>Cl.2=328.48 | Cl.1=41.80<br>Cl.2= 31.19 | Cl.1=1137<br>.6 Cl.2=<br>1101 | 77.63 |
|  | p | **<0.001** | **<0.001** | **Cl.1=<0.001**<br>**Cl.2=<0.001** | **Cl.1=0.0021**<br>**Cl.2=0.0027** | **<0.001** | **<0.001** |
|  | CI | 0.0002 | 0.0002 | 0.0002 | Cl.1=<0.001<br>Cl.2=0.001 | 0.0002 | 0.0003 |

Nonetheless, these results show that gamma ERS amplitude is modulated by target distance and further indicate that such modulation might reflect some kinematic features planned before movement onset.

### Movement-related gamma ERS and phase connectivity predicts peak velocity amplitude and timing

As both kinematic parameters and gamma ERS showed strong dependence on target extent, we explored the relationship between the amplitude and phase-synchronization of gamma oscillations and kinematic features by fitting linear regression models.

To model the relationship between gamma ERS amplitude and kinematics, we first extracted the z-transformed power of all the gamma bands from the two significant clusters of electrodes and used it as a predictor of peak velocity amplitude and movement time. We found that gamma amplitude during the planning and movement windows were significant predictors of movement time and peak velocity amplitude for all the tested gamma frequency bands (Table 2a). We also modeled the amplitude of gamma phase-synchronization in the network that was activated during the movement window. In this case, we extracted and z-transformed the wPLI values from the network edges that showed greater broad gamma and high gamma functional connectivity during movement execution compared to planning. Linear regression models showed that high gamma phase wPLI values in the highlighted sub-network significantly predicted peak velocity amplitude and movement time (Table 2b). Interestingly, no significant correlations were observed for the whole gamma band (Table 5b), thus suggesting that peak velocity and movement time might specifically depend on the amount of coherent high gamma activity in the highlighted occipital and contralateral parieto-frontal network. Both for gamma amplitude and wPLI, greater values were associated with higher peak velocity amplitude and shorter movement times, in line with the evidence supporting the prokinetic role of gamma oscillations.

**Table 2.** Linear regression models on the relationship between gamma amplitude and phase-coherence and kinematic features. a. Linear regression modelling of the z-transformed gamma amplitude and peak velocity (PV) amplitude and movement time (MT) in the time-windows corresponding to movement planning (−152 to −52ms) and execution (52 to 500ms). b. debiased WPLI values were extracted from the network showing greater phase coherence during movement execution compared to planning and used to predict PV and MT. F: F-statistic vs. constant model; Beta: coefficient estimate for the term in the model; SE: Standard Error of the coefficient; t: t statistic testing the null hypothesis that the corresponding coefficient is zero, p: p value; R^2^adj: adjusted coefficient of determination (df=60). In bold are reported the significant p values (alpha=0.05).

| **Peak velocity** | **Time window** | **F** | **Beta** | **SE** | **t** | **p** | **R<sup>2</sup>adj** |
| --- | --- | --- | --- | --- | --- | --- | --- |
| All Gamma<br>(25.5-80 Hz) | planning | 22.8 | 0.52 | 0.11 | 4.78 | **0.00012** | 0.26 |
|  | execution | 17.9 | 0.48 | 0.11 | 4.23 | **0.0008** | 0.22 |
| Medium Gamma<br>(40.5-55 Hz) | planning | 23.2 | 0.53 | 0.11 | 4.81 | **0.0001** | 0.27 |
|  | execution | 18.6 | 0.49 | 0.11 | 4.31 | **0.00062** | 0.22 |
| High Gamma<br>(55.5-80 Hz) | planning | 17 | 0.47 | 0.114 | 4.13 | **0.00011** | 0.21 |
|  | execution | 13.6 | 0.43 | 0.12 | 3.68 | **0.0005** | 0.17 |

| **Movement time** | **Time window** | **F** | **Beta** | **SE** | **t** | **p** | **R<sup>2</sup>adj** |
| --- | --- | --- | --- | --- | --- | --- | --- |
| All Gamma<br>(25.5-80 Hz) | planning | 15.1 | -21.26 | 5.47 | -3.89 | **0.00025** | 0.19 |
|  | execution | 12 | -0.41 | 0.12 | -3.46 | **0.001** | 0.15 |
| Medium Gamma<br>(40.5-55 Hz) | planning | 14.9 | -21.13 | 5.48 | -3.86 | **0.0003** | 0.18 |
|  | execution | 12.8 | -19.88 | 5.55 | -3.58 | **0.0007** | 0.16 |
| High Gamma<br>(55.5-80 Hz) | planning | 13.7 | -20.46 | 5.52 | -3.71 | **0.0005** | 0.17 |
|  | execution | 10.8 | -18.49 | 5.63 | -3.28 | **0.0017** | 0.14 |

**Table 2. Linear regression models on the relationship between gamma amplitude and phase-coherence and kinematic features.**
|  |  | **F** | **Beta** | **SE** | **t** | **p** | **R<sup>2</sup>adj** |
| --- | --- | --- | --- | --- | --- | --- | --- |
| WPLI debiased<br>(25.5-80 Hz) | MT | 3.42 | -0.71 | 0.38 | -1.85 | 0.069 | 0.07 |
|  | PV | 2.83 | 0.37 | 0.22 | 1.68 | 0.098 | 0.03 |
| WPLI debiased<br>(55.5-80 Hz) | MT | 11 | -0.496 | 0.149 | -3.32 | **0.0015** | 0.14 |
|  | PV | 14.2 | 0.366 | 0.097 | 3.77 | **0.0004** | 0.18 |

## Discussion

The present study confirms previous findings that movement-related cortical gamma synchronization during movement execution peaks around 70 Hz is parametrically modulated by movement extent. It further shows that such a parametrical modulation is present during movement planning with gamma peak occurring around 54 Hz. In addition, we found that gamma activity during the planning and execution of reaching movements displays different topographies. Finally, we showed that the amplitude of the movement-related gamma synchronization and its connectivity pattern are significant predictors of peak velocity amplitude and movement time.

It has been proposed that neocortical gamma oscillatory activity is central to sensorimotor and cognitive functions, specifically for information processing through the integration of neuronal assemblies’ activity, enabling a transfer of information across connected regions. Movement-related changes in the amplitude of gamma oscillations have been observed with ECoG in a variety of motor tasks specifically linked to some kinematic features such as movement trajectory (Talakoub et al., 2017), velocity (Bundy et al., 2016; Lofredi et al., 2018; Wang et al., 2017) and force (Flint et al., 2014; Jiang et al., 2020). A notable finding of the present study is that cortical gamma synchronization occurs during both planning and execution of the reaching movements and is parametrically modulated by movement extent. Increased oscillatory gamma activity, mostly in the medium (40.5-55 Hz) and high (55.5-80 Hz) frequency ranges, was visible immediately before and after movement onset, albeit with different topographies that may reflect different functions carried out during movement planning and execution. Interestingly, during movement planning, the first burst of gamma ERS was maximally expressed over the centro-parietal region, whereas the two gamma ERS bursts corresponding to the out- and back-movements were mostly visible over the parieto-occipital region.

The occurrence of gamma synchronization during the planning phase has been reported in previous studies (Donner et al., 2009; Gaetz et al., 2013; Gunduz et al., 2016; Ryun et al., 2017) and supports the idea that gamma ERS might be associated with top-down mechanisms engaged for movement preparation and, thus, with the creation of an efferent copy of the upcoming movement. Indeed, our task instructions (that is, to perform fast, uncorrected, out- and back-movements to targets appearing in an unpredictable order) mostly engaged feed-forward specification of a force-scaling factor according to movement extent (Ghez and Gordon, 1987; Gordon et al., 1994b, 1994a). Planning of movement distance between the starting hand position and the target requires proper jerk, acceleration and velocity scaling prior to movement onset (Gielen et al., 1985), a process that engages several regions including the primary sensorimotor cortices, supplementary motor area and the basal ganglia (Bell et al., 1994; Ghilardi et al., 2000; Tankus et al., 2009; Turner et al., 1998). Accordingly, the topographical distribution of gamma ERS before movement onset and its correlation with movement features suggest that the first burst of gamma ERS might reflect the engagement of sensorimotor neuronal assemblies appropriate to the characteristics of the upcoming movement. Indeed, our regression analyses confirmed that the amplitude of the pre-movement ERS positively predicts the amplitude of the peak velocity and, conversely, negatively predicts the movement duration.

Although the observed scaling of the pre-movement gamma ERS supports our interpretation of its role in movement planning, we cannot completely exclude that gamma ERS prior to movement onset might arise from subtle muscle contractions occurring before the motor act that can be captured only by comprehensive electromyography (EMG) recordings. Therefore, future analyses should combine EEG, EMG and kinematic recordings in order to clarify the relationship between pre-movement cortical gamma oscillations and muscle activation.

After the initial gamma ERS, gamma oscillatory activity visibly increased after movement onset and strikingly tracked the temporal profile of movements’ velocity. Compared to the initial burst that peaked around 54 Hz, this movement-related synchronous activity was most prominent in the high gamma range, in agreement with different studies (Cheyne and Ferrari, 2013; Joundi et al., 2012; Muthukumaraswamy, 2013; Pfurtscheller et al., 2003). Although gamma ERS was visible on the majority of the scalp sensors, our finding of greater synchronization over the posterior regions might reflect the engagement of the associative posterior parietal area, which is thought to integrate efferent motor signals with visual and proprioceptive information (Andersen et al., 1997; Bernier and Grafton, 2010; Elliott et al., 2017; Galletti et al., 2003; Hyvärinen and Poranen, 1974; Mountcastle et al., 1975; Shadmehr et al., 2010; Wolpert et al., 1998), as well as to encode and integrate visual and proprioceptive information during movement execution (Buneo and Andersen, 2012). This hypothesis is in line with the results of our phase synchronization-based connectivity analyses: compared to the planning window, during movement execution, we observed greater wPLI connectivity in the high gamma band between the electrodes located over the right parieto-occipital region and those over the bilateral fronto-central region. The same connectivity analysis on the broad gamma band displayed a more lateralized connectivity pattern between the left centro-parietal and fronto-central regions with a minor contribution of the parieto-occipital areas. Importantly, our regression analyses further showed that the network phase-synchronization index was a significant predictor of peak velocity and movement time in high gamma, but not that in the broad gamma range.

Altogether, we may speculate that movement-related ERS in the high gamma range during motor execution could reflect the specific activity of the dorso-medial stream, integrating online afferent sensorimotor and visual input and sending real-time feedback to premotor and motor regions to control the ongoing movement. Moreover, the observed positive linear relation between network connectivity and ERS amplitude and peak velocity might reflect either the amount of neural assemblies recruited to produce more force to increase speed (Muthukumaraswamy, 2010) or the amount of coordinative control needed to sustain the incoming afferent information about the ongoing movement.

In conclusion, this study provides an extensive characterization of scalp recorded movement-related gamma synchronization during planar reaching movements. Gamma ERS during movement planning and execution, as well as increased parieto-occipital and fronto-central connectivity during movement execution reliably predict the specific movement features.

## Methods

### Subjects and experimental design

We enrolled sixty-four right-handed healthy subjects (mean 23.9±4.5 years, 39 women) with normal or corrected vision, and without known disorders of the nervous system. This investigation was approved by the CUNY University Integrated Institutional Review Board (UI-IRB) and performed in accordance with the ethical principles of the Declaration of Helsinki and its subsequent amendments. Each participant signed an IRB-approved informed consent form before completing the experiment.

High-density EEG was recorded with the 256-channel HydroCel Geodesic Sensor Net (Electrical Geodesics Inc., Eugene, OR) while the participants completed a single block of 96 planar reaching movements (*mov* test).

### mov test

Participants performed 96 out-and-back reaching movements with their right dominant hand by moving a cursor on a digitizing tablet to targets appearing on a computer monitor. The targets were circles randomly presented every 3 s at three distances (4, 7 and 10 cm; radius: 0.5, 1, 1.25 cm, respectively) in eight directions (45° separation) (Figure 1a). The central starting point and the cursor position were always visible. Instructions were: to move the cursor as soon as possible without corrections, but only after the target presentation, with movements as fast and as accurate as possible, with overlapping strokes with fast reversals in the target circle.

### Kinematic data recording and analyses

The (x,y) coordinates of each trajectory were recorded with a custom-designed software and analyzed using an ad-hoc MATLAB-based pipeline. First, we filtered the coordinates with a Butterworth filter and then computed the first, second and third derivative of the trajectory to obtain velocity, acceleration and jerk for all the movements.

As detailed in previous publications (Ghilardi et al., 2000; Nelson et al., 2017; Perfetti et al., 2011), we computed several measures for each movement. In this study, we focused on: reaction time (i.e., the time from target appearance to movement onset), movement time (i.e., the duration of the outgoing movement), total movement time (i.e., the duration of the out and back movement), amplitude of peak velocity of the out-going segment (Figure 1a). Movements with any of such measures outside two standard deviations and those rejected from EEG preprocessing were excluded from EEG analyses.

#### Statistical analysis

Following Kolmogorov-Smirnov and Shapiro-Wilk normality tests on standardized residuals of each kinematic parameter, two participants were excluded from the analyses. Therefore, analyses of kinematic data were performed on the movements of 62 participants. Repeated measures ANOVA were run on reaction time, movement time, peak velocity amplitude and timing, and movement extent, with target distance (short, medium and long) as within-subjects factor. Violations of sphericity assumption were Greenhouse-Geisser-corrected and significant main effects (p < .05) were followed by Bonferroni-corrected pairwise comparisons. Further, we computed linear mixed-effect regression models using the MATLAB function *fitlme* to unveil the specific contribution of peak velocity and movement time on movement extent. Linear mixed-effects regression analysis is a versatile extension of simple linear regression models with excellent statistical power, as it allows the estimate of both fixed and random effects. Importantly, as mixed-effect models fit an intercept and/or a slope for each random-effect, they address the problem of the non-independency of the data (i.e. the inclusion of multiple trials per subject), while respecting between-subject variability. For each subject, we included all the available trials and set peak velocity and movement time as fixed-effect factors; the 62 participants were instead included as a random-effect factor. To best account for between-subjects variability, we tested the fit of a model including either an individual intercept or intercept and slope for each participant. To ascertain whether and which fitted model provided the best fit to the data, the following metrics were assessed: adjusted R^2^, Bayesian Information Criterion (BIC, an index used in Bayesian statistics to select among two or more models), and the Theoretical Likelihood Ratio Test (TLRT), which is commonly used to compare the goodness of fit of two statistical models. Specifically, the TLRT compared the model with random intercept and slope and the one with random intercept by computing their likelihood ratio test under the Chi-square distribution.

Importantly, even if visual inspection of residual plots did not reveal heteroscedasticity and deviations from normality, all the included values were z-transformed to obtain standardized estimates from the regression models.

### EEG recording and analyses

High-density (HD) EEG data were acquired using a 256-channel HydroCel Geodesic Sensor Net (Electrical Geodesic Inc.) with a Net Amp 300 amplifier (250 Hz sampling rate, online reference electrode: Cz) and Net Station software (version 5.0). Sampling frequency was 250 Hz and channel impedances were maintained below 50 kΩ throughout the recording to preserve a good signal-to-noise ratio.

All recorded data were preprocessed using the public Matlab toolbox EEGLAB version 13.6.5b (v.2016b) (Delorme and Makeig, 2004). The continuous signal was first filtered using a Finite Impulse Response Filter (FIR) between 1 and 80 Hz and Notch filtered at 60 Hz (59-61 Hz). Then, the signal was divided in 4-s epochs centered on target onset (−1 to 3 s) and visually inspected to remove sporadic artifacts and channels with poor signal quality.

Independent Component Analysis (ICA) with Principal Component Analysis (PCA)-based dimension reduction (max 108 components) was run to delete stereotypical artifacts, such as eye blinks, muscular activity and heartbeat. After a visual inspection of the power spectral density, topographical maps and time course of each estimated component, we retained an average of 16.05±6.27 components per subject. Channels previously removed due to bad signal quality were reconstructed using spherical spline interpolation, whereas those located on the cheeks and neck were removed. Re-reference to overall signal average was finally applied on the resulting 180 channels.

All the subsequent analyses were carried out using custom data analysis scripts with the MATLAB-based Fieldtrip Toolbox (Oostenveld et al., 2011).

Importantly, to avoid ambiguous effects from improperly executed movements, after the preprocessing, we discarded epochs representing movements whose kinematic parameters exceeded two SD.

After trial rejection, the average number of trials per subject was 76.2±13.2 SD, with a similar number of trials for each target distance (short: 25.6±4.6, medium: 26.2±5.9, long: 24.4±5.2). Data were then time-locked to movement onset (−1 to 2.5 s). Time-frequency representations (1-80 Hz) were computed using Complex Morlet Wavelets (0.5 Hz bins, 10 cycles). Each trial was baseline corrected by subtracting and dividing the average signal of the entire time-window of all trials.

#### EEG statistical analyses

##### Movement-related gamma oscillatory dynamics

EEG spectral and time-frequency analyses were run using the non-parametric cluster-based permutation procedure implemented in Fieldtrip (Maris and Oostenveld, 2007). Briefly, t or F statistics was first computed for each data point using a critical alpha of 0.001 and a minimum number of four significant neighboring electrodes to form a cluster. Cluster-level statistics was then run using the sum of the t/F values within each cluster of electrodes; the largest statistic value from the cluster-level analysis was then compared with a distribution of maximum cluster values obtained with 10000 permutations (Monte Carlo method, alpha=0.0005).

In order to characterize gamma oscillatory activity (25.5-80 Hz) during the planning and execution phases of the reaching movements, we first ran non-parametric permutation statistics and compared the broad gamma band time-course activity with the average spectral power (paired t-statistics, time window: − 500 to 2400 ms in 24 ms-time bins). Significant consecutive time-windows were then used to run second-level analyses to summarize the topography of different sub-bands of gamma (gamma: 25.5-80 Hz, low gamma: 25.5-40 Hz, medium gamma: 40.5-55 Hz, high gamma: 55.5-80 Hz). With the resulting clusters of electrodes (ROIs), time-windows and gamma frequency bands, we then explored whether target distance affected gamma amplitude. Thus, cluster-based permutation statistics were run with target-distance as within-subjects factor (repeated-measure ANOVA, planning time-window: −152 to −52 ms; movement execution time-window: 52 to 500 ms), along with the correspondent permutation-based post-hoc tests.

##### Movement-related gamma peak frequency analysis

To investigate whether the different phases of the reaching movements would be also characterized by spectral differences within the gamma range, we extracted for each subject the peak frequency value during the two significant time windows (planning and execution, see above) and a post-movement control time-window (1500-2000 ms after movement onset), when movements were certainly completed. Due to the distribution of the independent variable, normality assumption could not be satisfied (Shapiro-Wilk test on both data and residuals resulted in p<0.05). Therefore, non-parametric related-samples Friedman’s test was run to check for peak frequency differences in the three time-windows. Post-hoc pairwise comparisons were obtained with Dunn’s test and Bonferroni correction for multiple-comparisons (alpha=0.05).

##### EEG Functional Connectivity Analysis

To characterize gamma functional connectivity during movement planning and execution, we computed the squared weighted phase lag index (wPLI), a measure of phase-synchronization implemented in Fieldtrip as the debiased wPLI (Vinck et al., 2011). The wPLI derives from Stam’s Phase Lag Index (Stam et al., 2007), as it introduces a phase-difference weighting normalization with the imaginary component of the cross-spectrum (Nolte et al., 2004), thus improving robustness to noise. Also, the wPLI has the advantage of not being spuriously affected by the volume conduction of independent sources to different sensors or by a common reference, and shows increased statistical power to highlight true changes in phase-synchronization. First, we computed the power and cross-spectrum during the planning and execution time-windows using Complex Morlet wavelets (0.5 Hz bins, width=7) for two gamma frequency-bands (all gamma: 25.5-80 Hz, high-gamma: 55.5-80 Hz) and then estimated the debiased wPLI across all channel-pairs. The result of this process was a weighted network (for each time-window and each frequency band) represented as a 180 × 180 adjacency matrix C=[c_ij_], where each node is represented by a given electrode and each edge as the node-wise functional connectivity estimated by the wPLI. The functional connectivity matrices were exported to be statistically analyzed with Network Based Statistics (Zalesky et al., 2010).

##### Movement-related gamma functional connectivity

The Network Based Statistics (NBS) method was applied to detect eventual wPLI-based functional connectivity subnetworks differences during the movement and planning windows. This approach permits multiple hypothesis testing at the level of interconnected sub-networks, while controlling the family-wise error when performing analyses associated with a particular effect or contrast of interest (Zalesky et al., 2010).

NBS performs a mass univariate testing in order to identify the connections exceeding a test statistic threshold belonging to a given connected component. Then, a corrected p-value is computed for each component using the null distribution of maximal connected component size, which is empirically derived via a nonparametric permutation method. Here, a test with 5000 random permutations was performed to compute statistical significance for the identified network component. Finally, the hypothesis test is performed for the empirically determined components by comparing their extent with the proportion of permutations yielding a component with equal or greater size, correcting for the family-wise error rate at cluster level with p < 0.05. Here, a primary threshold t-score of 4.8 was chosen.

The results of the NBS procedure are presented as three-dimensional graph visualizations, which represented p<0.05 connection pairs surviving multiple comparison correction. The resulting functional connectivity values averaged over the significant sub-networks were then correlated with peak velocity and movement time to unveil a possible relation between gamma phase-synchronization in the identified sub-network and kinematic performance (see below).

##### Linear regression model on EEG data

Finally, in order to unveil a possible relationship between the observed changes in gamma oscillatory activity and the peak velocity amplitude and movement time, we fitted linear regression models to predict the kinematic parameters based on the gamma power during the planning and execution time-windows and the network connectivity values. Specifically, for the regression models with gamma amplitude as the predictor, we entered the average gamma power for the significant time-window (planning and execution), gamma sub-band and electrodes as independent variable and peak velocity and movement as dependent variables.

For the connectivity regression analyses, we extracted the average debiased wPLI value for the significant sub-network differences between planning and execution and used to predict peak velocity and movement time.

As reported for the kinematic analysis, in order to satisfy normality assumption, the data were z-transformed before model fitting. Further, two subjects were removed from all regression analyses as their kinematic average values were more than 2 SD lower than the mean performance.

## Acknowledgments

This work was supported by NIH P01 NS083514 (GT, CC, MFG) and by DOD W81XWH-19-1-0810 (AQ, MFG). Kinematic data were collected with custom-designed software, MotorTaskManager, produced by ETT s.r.l.. We thank Drs. Aaron Bruce Nelson and Serena Ricci for data collection; Martina Bossini Baroggi, Giorgia Marchesi and Giulia Aurora Albanese for the development of the kinematic analysis program (Marky) and Shaina George, Nancy Lin, Bunmi Aruleba and Henry Chen that helped in the analysis of the kinematic data.

## Author Contributions Statement

Study conception and design: MFG, ET, AQ. Data collection and analyses: ET, MFG, FF, AC, CC. Interpretation of the results: ET, MFG, AC, AQ, FF. Manuscript drafting: ET, MFG, AQ, FF, AC. Manuscript revising: ET, FF, MFG, AQ.

All the authors agreed to be accountable for all aspects of the work in ensuring that questions related to the accuracy or integrity of any part of the work are appropriately investigated and resolved.

