## Supplemental material for "Modulation of gamma spectral amplitude and connectivity during reaching predicts peak velocity and movement duration"

| | | RT | MT | PV Time | PV Amplitude | Movement Extent |
|---|---|---|---|---|---|---|
| **Distance** | **df** | 2 122.00 | 1.30 79.03 | 1.58 96.53 | 1.17 71.46 | 1.75 106.89 |
| | **F** | 24.26 | 285.52 | 305.80 | 868.72 | 5496.98 |
| | **p** | **< 0.001** | **< 0.001** | **< 0.001** | **< 0.001** | **< 0.001** |
| | **η²p** | 0.29 | 0.82 | 0.83 | 0.93 | 0.99 |
| **Short vs Medium** | **μ diff** | 7.52 | -22.44 | -12.65 | -17.85 | -3.07 |
| | **SE** | 1.19 | 1.42 | 0.77 | 0.57 | 0.05 |
| | **p** | **< 0.001** | **< 0.001** | **< 0.001** | **< 0.001** | **< 0.001** |
| | **CI** | 4.60 10.43 | -25.94 -18.94 | -14.56 -10.75 | -19.26 -16.43 | -3.18 -2.95 |
| **Short vs Long** | **μ diff** | 7.21 | -37.78 | -21.09 | -32.15 | -5.75 |
| | **SE** | 1.22 | 2.08 | 1.05 | 1.05 | 0.06 |
| | **p** | **< 0.001** | **< 0.001** | **< 0.001** | **< 0.001** | **< 0.001** |
| | **CI** | 4.22 10.21 | -42.91 -32.65 | -23.69 -18.49 | -34.73 -29.57 | -5.91 -5.59 |
| **Medium vs Long** | **μ diff** | -0.30 | -15.34 | -8.43 | -14.31 | -2.68 |
| | **SE** | 1.26 | 1.11 | 0.71 | 0.60 | 0.05 |
| | **p** | 1.00 | **< 0.001** | **< 0.001** | **< 0.001** | **< 0.001** |
| | **CI** | -3.41 2.80 | -18.07 -12.61 | -10.18 -6.68 | -15.79 -12.82 | -2.81 -2.55 |

**Supplementary Table 1**. Results of repeated measure ANOVAs on the effect of target distance on the behavioral indices. RT: Reaction Time; MT: Movement Time; PV: Peak Velocity; μ diff: mean difference; SE: Standard Error of the mean; CI: Confidence Interval; η²p: partial eta squared.

| | FE | Beta | CI | | t | $p_t$ | $R^2$adj | BIC | TLRT | $p_{TLRT}$ |
|---|---|---|---|---|---|---|---|---|---|---|
| **Model with random intercept** | PV | 0.99 | 0.98 | 1.0 | 204 | **< 0.0001** | 0.92 | 1792.7 | - | - |
| | MT | 0.63 | 0.62 | 0.65 | 93 | **< 0.0001** | | | | |
| **Model with random intercept & slope** | PV | 1.06 | 1.01 | 1.1 | 2193 | **< 0.0001** | 0.95 | -174.5 | 2009 | **< 0.0001** |
| | MT | 0.70 | 0.65 | 0.75 | 905 | **< 0.0001** | | | | |

**Supplementary Table 2.** Linear mixed-effects regression models on kinematic data. (Top) Mixed-effect regression model including a random intercept for each subject; (Bottom) Mixed-effect regression model including random intercept and slope. FE: Fixed-effects included in the models (PV: Peak Velocity amplitude; MT: Movement Time); Beta: Estimate of the slope for the FE; t= t test; CI: Confidence Interval of the FE estimate; BIC: Bayesian information criterion; $R^2$adj: adjusted coefficient of determination; TLRT: Theoretical Likelihood Ratio Test. $p_t$: p value associated to the t statistic testing the effect of the FE; $p_{TLRT:}$ p value resulting from the TLRT which tested which model showed a better fitting of the data. Significant results are reported in bold (alpha=0.05).
